# Viruses roam the wheat phyllosphere

**DOI:** 10.1101/2021.05.31.446456

**Authors:** Laura Milena Forero-Junco, Katrine Wacenius Skov Alanin, Amaru Miranda Djurhuus, Witold Kot, Alex Gobbi, Lars Hestbjerg Hansen

## Abstract

The phyllosphere comprises all the above-ground sections of plants. This niche is colonised by complex microbial communities, including algae, fungi, archaea, and bacteria^1–3^. They are known to induce plant growth and promote health^4,5^ or act as causative agents of plant diseases^6–8^. It is thought that the most abundant organisms are phyllobacteria^2^, with an estimate between 10^6^ to 10^7^ cells per cm^2^. Viruses are highly abundant across many environments, often outnumbering bacteria 10 to 1^9^. However, not much is known about their abundance and composition in the phyllosphere, a harsh environment for viruses due to environmental variability and high UV exposure. To investigate this niche in detail, we extracted, sequenced and analysed phyllosphere virome. Using leaf samples from winter wheat (*Triticum aestivum*), we identified a total of 876 viral populations (vOTUs), mostly belonging to the *Caudovirales* order. Most of these were predicted to be lytic. Remarkably, 810 of these viral populations correspond to new viral species with no matches to known sequences. We estimate a minimum of 2.0 x10^6^ viral particles per leaf. Overall, these findings suggest that the phyllosphere ecosystem harbours an abundant and active community of novel viruses that play essential roles in shaping this habitat.

---

Wheat has been a significant crop providing energy and essential nutrients for centuries^10^. The importance of understanding every aspect of the crop to optimise the yield is still relevant due to the growing human population and the threat of climate change^10^. The flag leaf is the ultimate leaf to emerge, and it is known to be a major producer of energy for the plant, and the properties of this leaf have been indicated to affect the crop yield^11^. The plant microbiome can heavily influence plant growth and is shaped by its multiple interactions with the environment and other biotic inhabitants^12^. Bacteriophages (phages), viruses of bacteria, are major players in shaping various microbial communities, including those in the phyllosphere^13–18^. However, little is known about their diversity and abundance in this habitat, and a phyllosphere virome has never been described before. Hence, we chose to examine the flag (FL) and penultimate (PL) leaves of the wheat plant *Triticum aestivum* cultivar Elixer to investigate the double-stranded DNA (dsDNA) and single-stranded DNA phyllovira (phyllosphere-associated viruses). In this study, we present the first two sequenced phyllosphere viromes. The wheat phyllovira were extracted by collecting 500 leaves per leaf type. The sampling was performed at Vridsloesemagle, Denmark, at the ripening/maturity stage. The DNA from the viral fraction was directly sequenced from the FL and PL. It has previously been shown that ssDNA phages and circular elements are selectively amplified by multiple displacement amplification (MDA)^19^. Therefore, two additional samples, FL_mda_ and PL_mda_, were sequenced after DNA amplification to get qualitative data on ssDNA viruses. In addition, the remaining microbial fraction was analysed by Illumina shotgun metagenomics (FL_mf_ and PL_mf_).

The viral fraction of the FL and PL samples resulted in 91.6 ng and 37.2 ng of DNA, respectively (Supplementary Table S1). The genome assemblies from the four viromes (FL, PL, FL_mda_, and PL_mda_) resulted in a total of 25,977 contigs, where 15,739 (60.6%) were putatively identified as viral, based on homology to previously identified viral sequences with VirSorter2^20^. They constitute 73.12 million bp (Mbp) which recruited the vast majority of the sequencing effort, as 84.2% of the quality-controlled reads mapped to said contigs. Non-putative viral contigs are either artefacts, bacterial remnants, or, likely, viral sequences with no homologs in viral databases. The viral contigs were clustered into 876 species-rank virus groups (95% sequence identity and 85% coverage), referred to as viral populations or viral operational taxonomical units (vOTUs)^21^. The vOTUs consist of high quality/complete viral genomes, or sequences of length >10 kbp^22^. The vOTU representatives recruit 72.2% of the sequencing effort, yet they only represent 27.1% of the assembled contig length and 3.37% of the total contig number (Fig. 1A). From the 876 vOTUs, 65 were identified as complete phage genomes, while 107 as high quality and 139 as medium quality (Fig. 1B). The genome length of the complete genomes ranges from 2,135 bp ssDNA phages up to 354,470 bp giant phages. Based on the amount of DNA extracted from the leaves and the average size of the complete genomes recovered from this study, we estimate a minimum of 6×10^4^ to 1.5×10^5^ viral particles per cm^2^ or in average 2.0 x10^6^ viral particles per leaf (Methods and Supplementary Table S1). By read mapping, we identified the presence of 400 vOTUs in the FL and 698 vOTUs in the PL. In contrast, we identified 183 and 368 vOTUs in the FL_mda_ and PL_mda_, respectively, showing an apparent loss of diversity in the MDA-amplified viromes (Fig. 1C). There were 66 vOTUs shared between the four viromes (FL, PL, FL_mda_, and PL_mda_), and we observed higher viral diversity in PL compared to FL. A total of 50 viral populations were unique to the amplified fractions, 5 of them annotated as high-quality ssDNA viruses (Fig. 1C).

**Figure 1.**
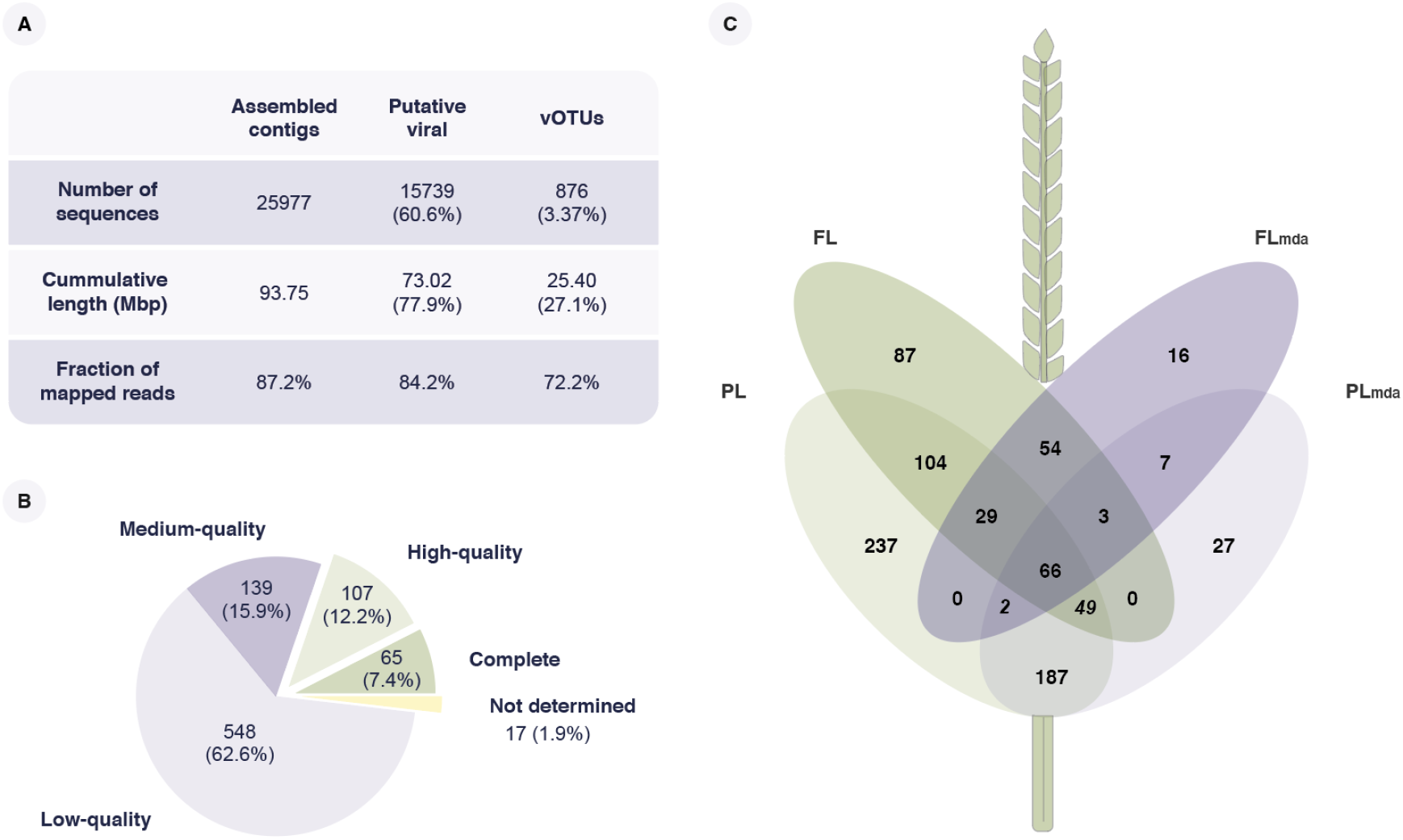
Recovered viral populations (vOTUs) from the four types of viromes (FL, PL, FL_mda_, and PL_mda_). **A**. Overview of contigs from the combined assembly, putative viral contigs and vOTU representatives, in terms of the number of sequences, cumulative contig length, and the fraction of mapped reads. **B**. Putative completeness of vOTU sequences. **C**. Venn Diagram of unique and overlapping vOTUs between FL, PL, FL_mda_, and PL_mda_.

From the microbial fraction shotgun data, the most dominating phyla observed on FL_mf_ and PL_mf_ samples are Actinobacteria, Bacteriodetes, Proteobacteria and Firmicutes (Fig. 2), consistent with previous findings of the wheat microbiome^23,24^. Both leaf types are dominated by the orders *Xanthomonadales* (33.14%) and *Pseudomonadales* (25.17%) (Fig. 2). When linking the bacterial composition of the two leaf types with the viral population, we found that 702 (80.6%) of vOTU matched a CRISPR spacer in the Dion database^25^. In addition, there were 27 vOTUs matching spacer sequences from the FL_mf_ and PL_mf_ microbial assemblies, 17 of which are assigned to bacterial hosts of the *Erwinaceae* family. Based on the CRISPR spacer sequences, we identified 78 vOTUs against *Pseudomonadaceae* and 21 vOTUs targeting the *Xanthomonadaceae*, the two most abundant bacterial families. Moreover, 154 vOTUs were identified to target the *Enterobacteriaceae* family, which only represents 0.78% of the relative abundance and is primarily found in the FL_mf_ and not in the PL_mf_ fraction.

**Figure 2.**
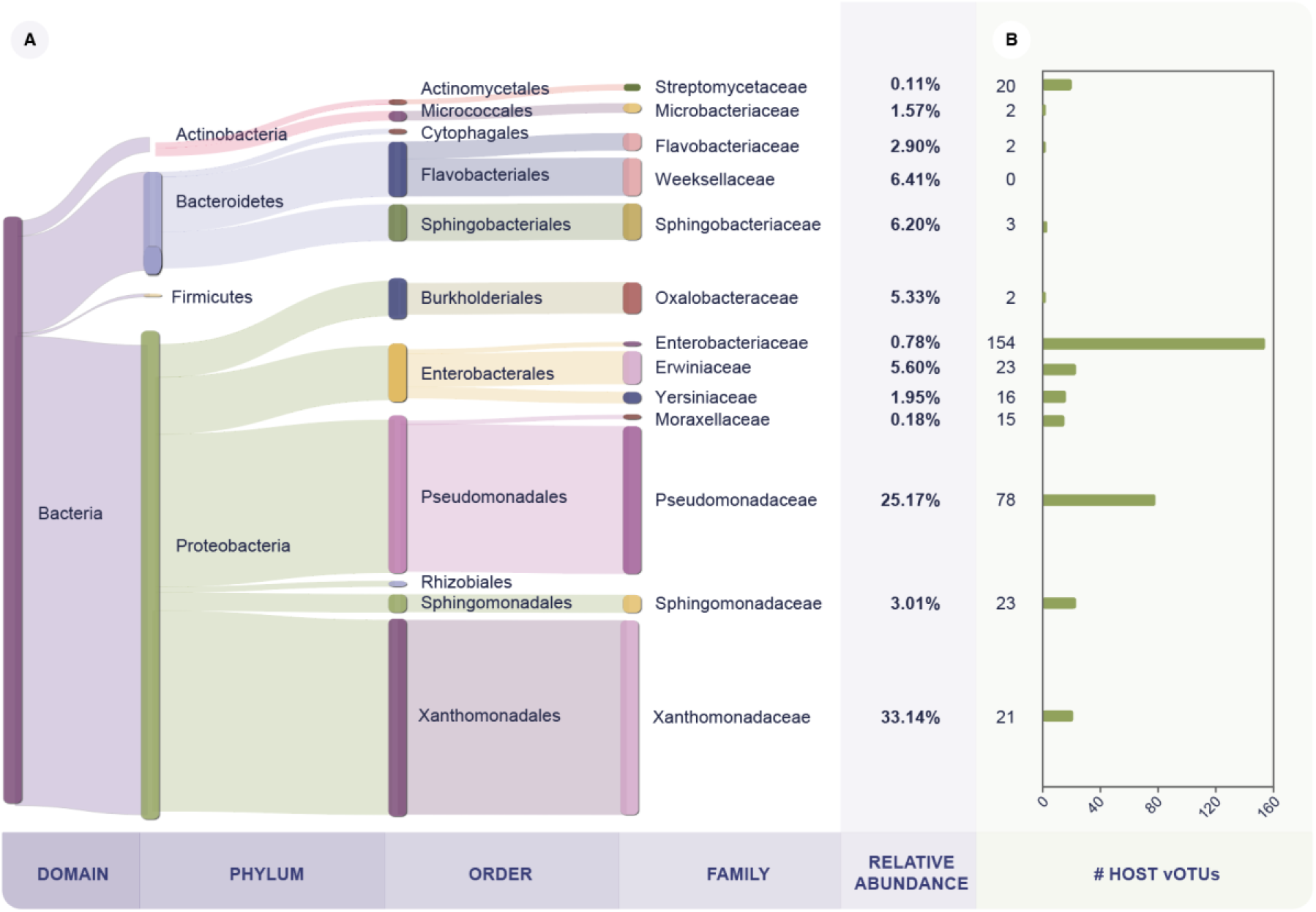
Microbial composition and viral-host taxonomy. **A.** Microbial relative abundance of the most abundant families. Taxonomy was assessed at read level from WGS samples (FL_mf_ and PL_mf_) using Kraken2. **B.** Number of phages linked to bacterial host at the family level. The viral hosts were predicted by matching CRISPR spacers from the Dion CRISPR spacer database^25^ to the wheat vOTUs sequences. *Moraxellaceae* and *Streptomicetaceae* are not abundant families in the microbial fraction but were included since they have ≥15 vOTU CRISPR matches.

We performed a detailed examination of the 10 most abundant vOTUs on each of the two leaves (Fig. 3). While most of these were abundant only in one of the two leaf pools, the vOTU classified as *Hamiltonella virus APSE-3* was the exception, with a relative abundance of 23.1 % and 3.1 % on the FL and PL respectively (Fig. 3A). Interestingly, as much as 36.8% of the FL reads map to the APSE-3 genome, compared to 4.5% for the PL. *Hamiltonella defensa* (*H. defensa*) is a facultative endosymbiont that lives in several species of aphids, constituting a tripartite insect-bacteria-phage symbiont system^26^. In this association, the phage integrates into the *H. defensa* genome and indirectly provides the aphid with protection from parasitic wasps^27^. The APSE phages identified in *H. defensa* genomes have been shown to have very conserved genomes, with most variations occurring in one highly variable region encoding a toxin cassette. Different putative toxins, including a Shiga-like toxin, a YD-repeat (RHS-repeat) toxin or a cytolethal distending toxin (CdtB), are encoded in this region, leading to the classification of distinct APSE variants encoding different putative toxins^27^. Interestingly, the abundant APSE type identified on both FL and PL seems to contain a YD-repeat putative toxin (Fig. 3B, S1), placing them in the APSE-3 group, which have been shown to confer the strongest defensive phenotype^27^. When mapping the reads obtained from FL and PL to the APSE-3 contig, there is a clear difference in coverage in the toxin-cassette encoding region, indicating distinct APSE populations, encoding different toxins on the two leaf types (Supplementary Fig S2). Furthermore, the low amino-acid identity in the YD-repeat toxin between the wheat APSE and other members of APSE-3 indicates that this could be considered a new variant, 3.2a (Fig. 3, Supplementary Fig. S1).

**Figure 3.**
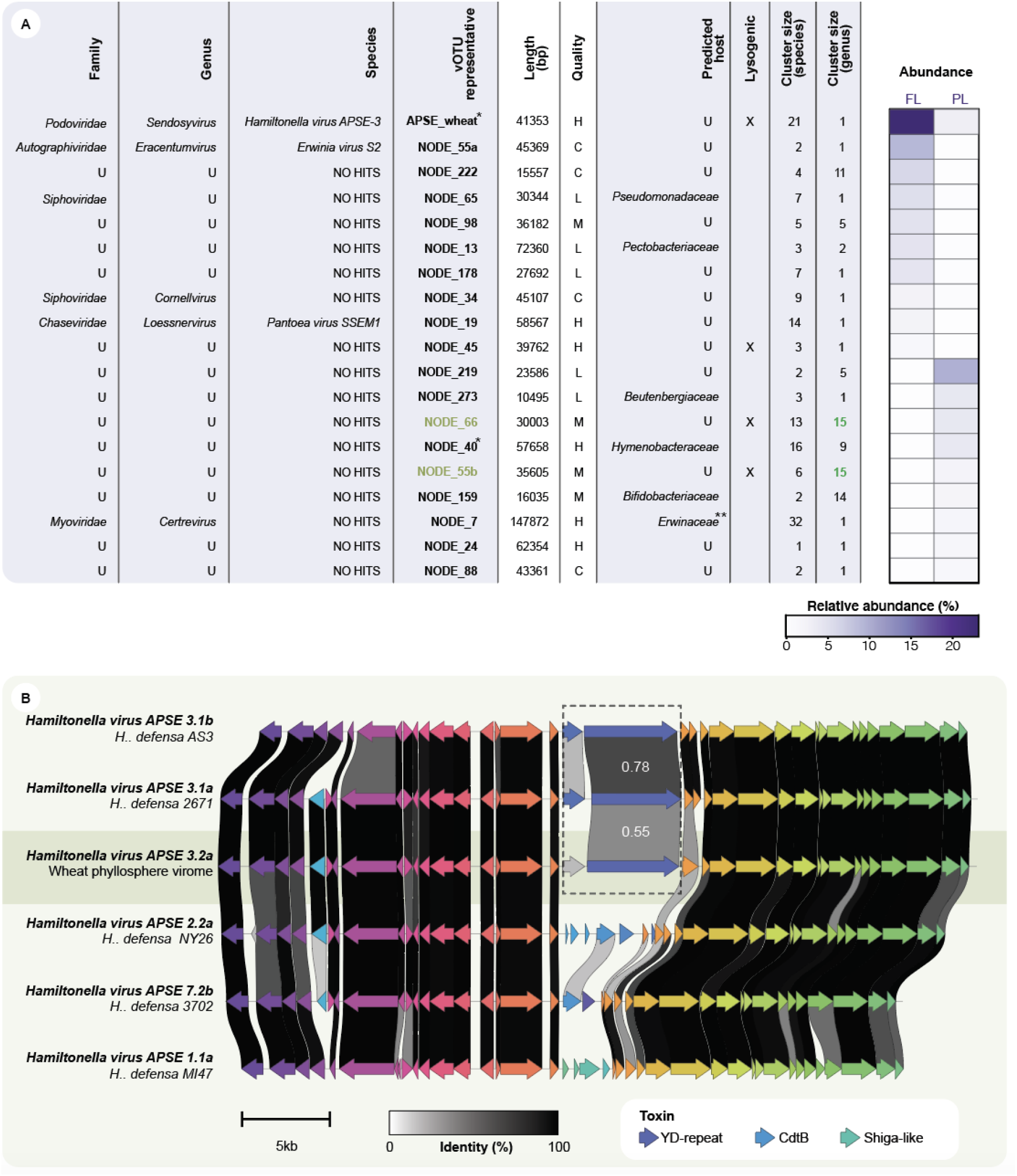
Description of the 10 most abundant phages on the FL and PL, and genome comparison of assembled APSE phage with APSE-3, APSE-2, APSE-7, and APSE-1 members. **A** Description of the 10 most abundant phages. Taxonomic assignment at family/genus level was performed with vConTACT2 and at species level using blastn (>85% coverage, >95% identity). This description includes the length of vOTUs representatives with genome quality assessed with CheckV and lysogeny with VIBRANT. Cluster size (species) indicates how many phyllosphere viral sequences group in a vOTU after clustering at 85% coverage, 95% identity. On the other hand, genus cluster size refers to how many vOTUs belong to the same genus cluster as per vConTACT2(C) Putative complete, (H) Putative high quality, (M) Putative medium quality, (L) Putative low-quality. (U) Unclassified). *Contigs assembled from a subsample, **Viral contig with a match to the WGS’s CRISPR spacer set. The abundance of the vOTUs, was calculated as the average number of reads overlapping each base, normalised by the number of reads on flag leaf (FL) and penultimate leaf (PL) samples **B**. Genome comparison of the wheat-phyllosphere assembled APSE phage and 5 previously described APSE virus types, with amino-acid identity indicated between the genomes. Open reading frames are coloured by identity.

Out of the most abundant vOTUs, three could be classified to the species level and six to the family level, indicating that many of these abundant vOTUs represent members, which could be unique to the wheat phyllosphere (Fig. 3A). Moreover, the 19 abundant vOTUs were assigned into 18 distinct genera, where only NODE_66 and NODE_55b grouped in a genus-level cluster, constituted by 15 viral species. Likewise, more than half (11 out of 19) of the abundant vOTUs are single members of a genus cluster, and these clusters can group up to 32 viral species (Fig. 3B). Interestingly, 4 out of 19 (15.7%) of the most abundant phages were classified as lysogenic, indicating that the extracted virome diversity is composed mainly of lytic phages, which is corroborated by only 134 out of 876 (15.3 %) of the vOTUs being identified as lysogenic. A UV-rich environment such as the phyllosphere is known to be detrimental to phages^28^. Therefore, it might have been expected to favour the protection provided to lysogenic phages by a host. However, the proportion of lysogenic to lytic phages has previously been linked with bacterial biomass, where lower microbial densities seem to favour ‘kill-the-winner’ dynamics resulting in an increased ratio of lytic phages^29^. While this first phyllosphere virome only represents a snapshot, it would be interesting to examine further if this trend is conserved across different spatiotemporal scales.

Lastly, the uniqueness of the viral sequences was investigated by comparing the vOTUs against previously identified viral sequences from the IMG/VR 3.0 database. This database comprises more than 2 million cultivated and uncultivated viral genomes, including reference viral genomes. At species level (95% identity, 85% coverage), only 57 (6.5%) of the vOTUs had significant hits to IMG/VR, indicating that 810 of the assembled vOTUs correspond to putatively new viral species. These 57 wheat-derived vOTUs match 271 sequences from the IMG/VR database. Then, a protein-sharing network was built to further explore and visualize the genomic relationship at the genus level to IMG/VR and RefSeq viral sequences (Fig. 4A). The network protein clustering via vConTACT2 allowed to link only 29 (3.3%) vOTUs to RefSeq genomes at the genus level. Moreover, it is possible to identify 17 groups in the network, composed only of wheat phyllovira (77 contigs). For instance, NODE_40, one of the most abundant species found in the PL, is part of one of these phyllosphere-only groups 29 other viral species (vOTUs). Based on the genus-level clustering, this group encompasses up to 13 distinct viral genera (Fig. 4B). This suggests that there are numerous novel, abundant and diverse groups of viruses in the phyllosphere virome that are likely unique to this environment.

**Figure 4.**
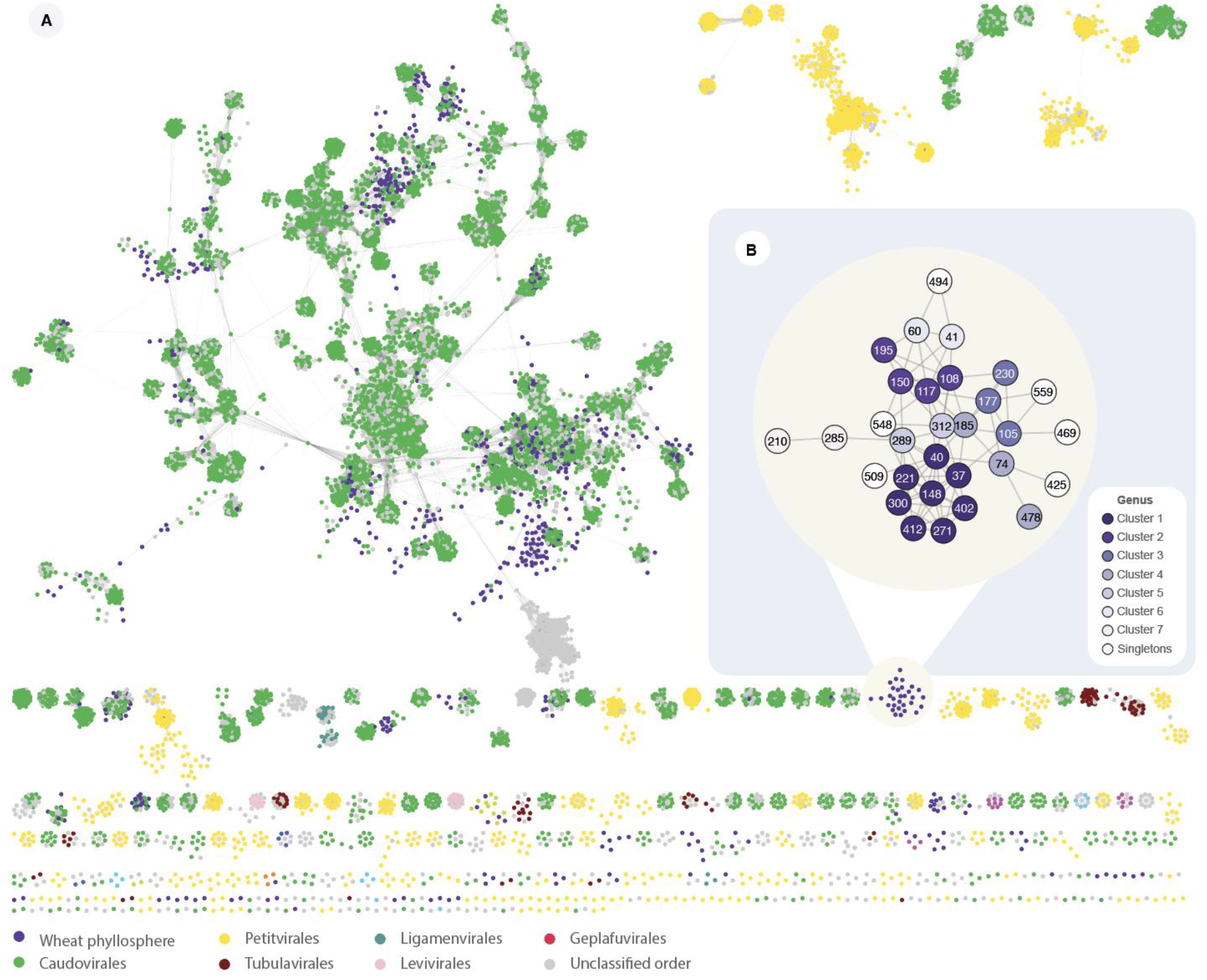
Viral protein sharing network created with vConTACT2 clustering wheat phyllosphere vOTUs and RefSeq and IMG/VR genomes. Each node represents a viral genome, and edges between two nodes indicate a shared protein profile. **A**. Colours represent the taxonomy of RefSeq sequences. **B** Zoom-in of one of the unique groups of phages found on the wheat phyllosphere, colours represent distinct vConTACT2 clusters.

In conclusion, we here present the first detailed description of a virome from a phyllosphere environment. We found a total of 876 viral species groups, or vOTUs, of which 810 were novel. We show that the wheat phyllosphere is composed of a diverse viral community with at least 2.0 x10^6^ viral particles per leaf. This is a very conservative estimate taking into consideration the DNA loss throughout the virome extraction. Furthermore, we demonstrate that phage microbial components interact with bacteria and should be taken seriously as a major player in shaping the phyllosphere microbiome. As such, they could dramatically affect plant health and crop yield. A previous study showed that the viral fraction of tomato leaves affected the bacterial composition, which is with high certainty the same case for the phyllosphere virome on the wheat leaves^18^. We show that two different communities of dsDNA and ssDNA phages inhabit the wheat flag and penultimate leaves, indicating that the viral composition is different along plant niches. This further supports that the phyllosphere is an underexplored biotope in bacteriophage ecology and underlines the need to explore phages in more diverse environments to expand the viral sequence information currently available in public databases. We observed that the most abundant phage on the FL was an APSE phage related to transkingdom interaction between phages, bacteria and aphid. These results represent an early beginning of a new chapter of phyllosphere biology, which should to address phage populations and not only bacteria and fungi.

## Methods

### Sample collection

A total of 500 flag leaves (FL) and 500 penultimate leaves (PL) from the wheat plant cultivar Elixir, were collected on June 8^th^, 2020 near Vridsloesemagle in Denmark, when the plants were in the ripening/maturity stage 11 (ref). Leaves were cut up and shaken at 200 rpm in 1 L SM-buffer (100 mM NaCl, 8 mM MgSO4, 50 mM Tris-Cl) for 1 hour at 4°C to extract the bacteriophages from the leaves into the buffer for concentration and DNA extraction. After manual removal of leaf cuts, the SM-buffer was centrifuged at 10,000 x g followed by sequential filtering with 5.0 μm and 0.45 μm (PVDF, Merck Millipore, Darmstadt, Germany) to remove plant cells, chloroplasts and larger microbes. Pellets from the centrifugation step were saved and used for microbial extraction.

### DNA extraction and amplification

Polyethylene glycol (PEG) precipitation was carried out to concentrate 1 L sample down to 10 mL SM-buffer containing the leaf virome as explained in the protocol ‘Harvesting, Concentration, and Dialysis of Phage’ from The Actinobacteriophage database (phagesdb.org/workflow/ last accessed September 6^th^ 2019). DNA extraction was carried out as previously explained^30^. Briefly, the two 10 mL concentrated virome samples (FL and PL) were treated with 83 U of DNAse I (A&A Biotechnology, Gdynia, Poland) for 1 hour at 37°C with CaCl_2_ + MgCl_2_ added in 10mM concentration to remove free DNA. Following with 18 U of proteinase K treatment (A&A Biotechnology, Gdynia, Poland) and 0.1% (v/v) sodium dodecyl sulfate (SDS) for 1 hour at 55°C to remove phage capsids and other proteins. The treatment was followed with Clean and Concentrator −5 (Zymo Research, Irvine, CA, US) DNA extraction. The two viromes were amplified using Cytiva illustra^TM^ Ready-To-Go^TM^ GenomiPhi V3 DNA Amplification Kit (Cytiva, Marlborough, MA, US). The bacterial pellets were collected from supernatant filtered through a 5.0 μm filter centrifuged for 10 min at 10,000 g. The pellets were re-suspended in the lyses buffer from the kit DNeasy Powerlyzer Powersoil Kit (QIAGEN, Hilden, Germany) and the DNA extracted according to the manufacturer’s procedure.

### Library prep of Virome and Bacterial WGS

The concentration and purity of the DNA from both the viral and microbial fraction were measured using the Qubit 2.0 fluorometer (Life Technologies, Carlsbad, CA, US) and NanoDrop 2000 spectrophotometer (Thermo Scientific, Waltham, MA, US), respectively. Libraries for amplified and non-amplified viromes, as well as the microbial fraction, was prepared according to manufacturer’s using the Nextera® XT DNA kit (Illumina, San Diego, CA, US) and sequenced as paired-end 2×150 bp reads on Illumina NextSeq500 platform using the Mid Output Kit v2 (300 cycles).

### Quality control and assembly

Raw sequencing quality was assessed with FastQC v.0.11.9^31^. Trimmomatic v.0.39^32^ was used for the removal of low-quality bases and adapters. Additionally, reads mapping to ΦX174 were removed using bbduk from BBtools suite v.38.86^33^. Subsets of each sample were created with 10, 20, 30, 40, 50, 60, 70, 80, and 90% of the reads. Samples were down normalised to a target coverage of ~100x, and error corrected with bbnorm (BBtools suite). Assembly was done with metaSPAdes v.3.14.0 ^34^ without the error correction stage. The resulting contigs were filtered to a minimum contig size of 1000 bp and 2x coverage. Contigs assembled using 100% of the reads were used for further analysis. Two full genomes were included from the additional assemblies, NODE_40, from the 80% subsampling of the PL, and APSE_wheat, from the 10% subsampling of PL_mda_.

### Viral Identification and viral clustering

VirSorter v.2.0^35^ was run on the assembled contigs to identify viral contigs. Then, the putative viral sequences were clustered into viral populations (species level) using stampede-clustergenomes script (https://bitbucket.org/MAVERICLab/stampede-clustergenomes/, <u>last accessed March 11^th^, 2019</u>). They were clustered at 95% nucleotide identity across 85% of the genome^21^, and the longest sequence was selected as a representative. The completeness of cluster representatives was predicted with CheckV v0.6.0 and vOTUs were defined as representative contigs >10kb or labelled as high quality/complete viral genomes^22^. Identification of lysogenic sequences was made with VIBRANT^36^.

### Abundance calculations

Quality controlled reads were mapped against the vOTUs using bbmap v.2.3.5^37^. Contig breadth coverage was calculated with genomecov from BEDtools v.2.29.2.0^38^. SAMtools v.1.10^39^ was used to calculate the number of mapped reads per contig, which was then normalized by contig length and sequencing depth. Contigs were required to have more than 75% of their genome covered to be considered present in a samples^22^. The fraction of mapped reads was calculated by calculating the percentage of quality-controlled reads mapped to assembled contigs, putative viral contigs, or filtered vOTUs. Coverage plots from BAM files were created with weeSAM version 1.5 (https://github.com/centre-for-virus-research/weeSAM, <u>last accessed September 22^nd^, 2020</u>)

### Bacterial and viral taxonomy

The bacterial composition of the FL and PL was assessed by classifying reads at the family level using Kraken v2.1.1 against the minikraken2_v28GB_201904 database. Viral taxonomy was assigned at species level using nucleotide BLAST against nr. Species were classified if there were hits with more than 85% coverage at 95% identity. Viral populations were classified at the genus level using vConTACT2 protein profile clustering against IMG/VR^40^ and viral RefSeq v94. vOTUs were aligned with nucleotide BLAST to the IMG/VR 3.0 database, and only contigs convering at least 40% of a vOTU, and with a genome ratio >0.8 were included in the network. Family level classification for bacterial and viral sequences was performed with MMSeqs2 v12.113e3 taxonomy module^41^. It was based on a translated search against RefSeq genomes, where the last common ancestor was calculated with the 2bLCA protocol, with a minimum query coverage of 30%.

### Host assignment

Two sets of CRISPR spacers were used to match viral sequences with putative host families. The first set was derived from identified spacers on the microbial contigs of the wheat phyllosphere, detected using MinCED (https://github.com/ctSkennerton/minced/, <u>last accessed September 22^nd^, 2020</u>). The other set was the Dion^25^ database containing >11 million spacers. Phage–host relationships were identified by matching viral sequences to CRISPR spacers using SpacePHARER^42^. In order to assign a host taxonomy, at family level, at least 80% of the CRISPR hits had to belong to the same family.

### Viral particle estimation

The following formula was used for computing the number of viral particles per leaf:

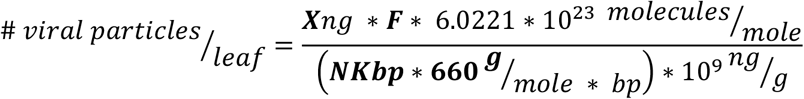

Where X corresponds at the amount of DNA extracted from the viral fraction (0.13 ng/leaf), F corresponds to the fraction of reads that map to viral contigs (0.84), and N corresponds to the average length of a bacteriophage genome (50Kbp). This is a total of 2.0 x10^6^ viral particles per leaf, which using a mean wheat leaf area of 20cm^2^, results in an estimate of 9.8 x10^6^ viral particles per cm^2^.

### Genome comparison

The wheat APSE genome was compared against 15 other previously described APSE genomes^27^. The detection of open reading frames and annotation of viral sequences were done with VIGA. Finally, gene clusters were calculated and visualised with clinker v0.0.20^43^. Based on the gene content observed in two variable regions of the APSE genomes (toxin cassette and DNA polymerase region), we propose the wheat phage to be classified as APSE type 3.2a.

## Data availability

The sequencing data used and described in this study has been submitted to NCBI under the Bioproject accession number PRJNA733924.

## Acknowledgements

This research was supported and funded by the Novo Nordisk Foundation with grant number NNF19SA0059348 (The MA*T*RIX) for LMFJ, AG and LHH. The research also received funding from the European Union’s Horizon 2020 research and innovation program under the Marie Sklodowska-Curie grant agreement No 801199 for LMFJ. The project also had partial support from Human Frontier Science Program with grant number HFSP-RGP0024/2018 for KWSA, WK and AMD. Special thanks to Svend Christensen and Jesper Cairo, from helping with the logistics of the the Vridsloesemagle sampling site, with affiliation to Department of Plant and Environmental Science, University of Copenhagen, Frederiksberg, Denmark.

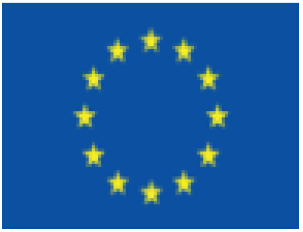

## Author contributions

LMFJ, LHH, and WK conceptualised the study. LMFJ, KWSA, and AMD coordinated and did the sampling of flag and penultimate leaves, and virome and DNA extraction. KWSA, AMD, and WK did the library preparation, amplification and sequencing of the virome DNA. AG and AMD extracted and prepared the bacterial fraction libraries. LMFJ did the bioinformatics analysis and the visualisation of all figures. LMFJ, AMD, KWSA, and AG wrote the manuscript, while WK and LHH supervised and added extensive edits throughout the manuscript.

