## Supplementary material for "Viruses roam the wheat phyllosphere": Table S1, Figure S1, and Figure S2

**Table S1**: **The DNA concentration of six different sample types.** Overview of DNA concentration from the microbial and virus fraction of both leaf types, as well as the DNA concentration of the amplified virus fraction.

| Sample | DNA conc. (ng/µl) | Sample size (µl) | Total amount of DNA | Raw sequencing reads |
|---|---|---|---|---|
| $FL_{mf}$ | 100 | 100 | 10 µg | 20347654 |
| $PL_{mf}$ | 8 | 100 | 800 ng | 20392188 |
| FL virome | 4.58 | 20 | 91.6 ng | 71400921 |
| PL virome | 1.86 | 20 | 37.2 ng | 72460435 |
| $FL_{mda}$ virome | 5.28 | 10 | 52.8 ng | 35481660 |
| $PL_{mda}$ virome | 1.22 | 10 | 12.2 ng | 24508019 |

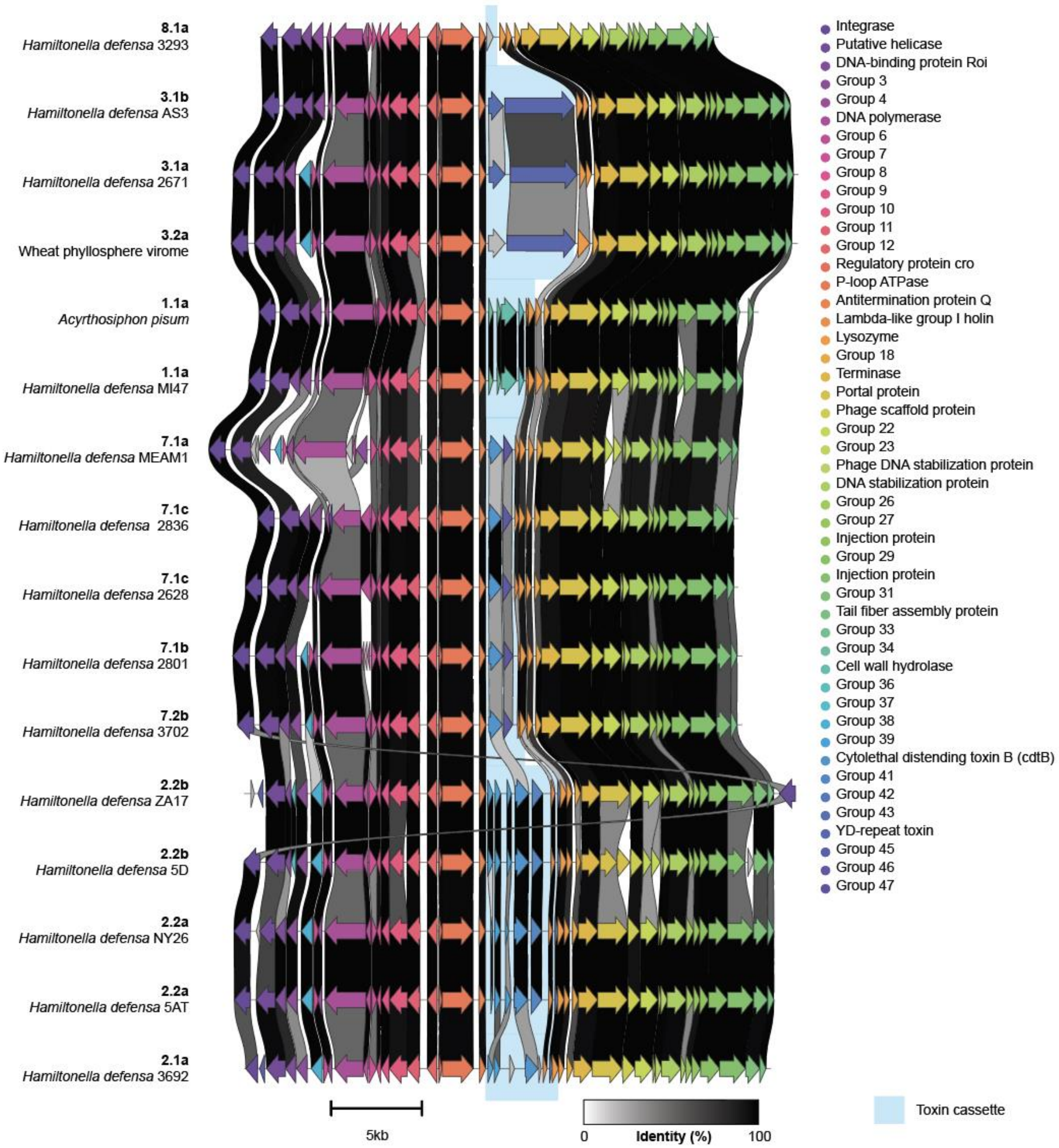


**Figure S1**. **New APSE-phage variant discovered in the wheat phyllosphere.** Genomic comparisons of representative variants in the APSE group at the amino acid level (clinker). The main variation between these members is observed in the region encoding a toxin cassette, carrying either shiga-like toxins, cytolethal distending toxins (CdtB), which describes the different variants. The wheat APSE vOTU resembles members of variant 3, encoding a YD-repeat toxin. However, with only 55% amino acid identity.

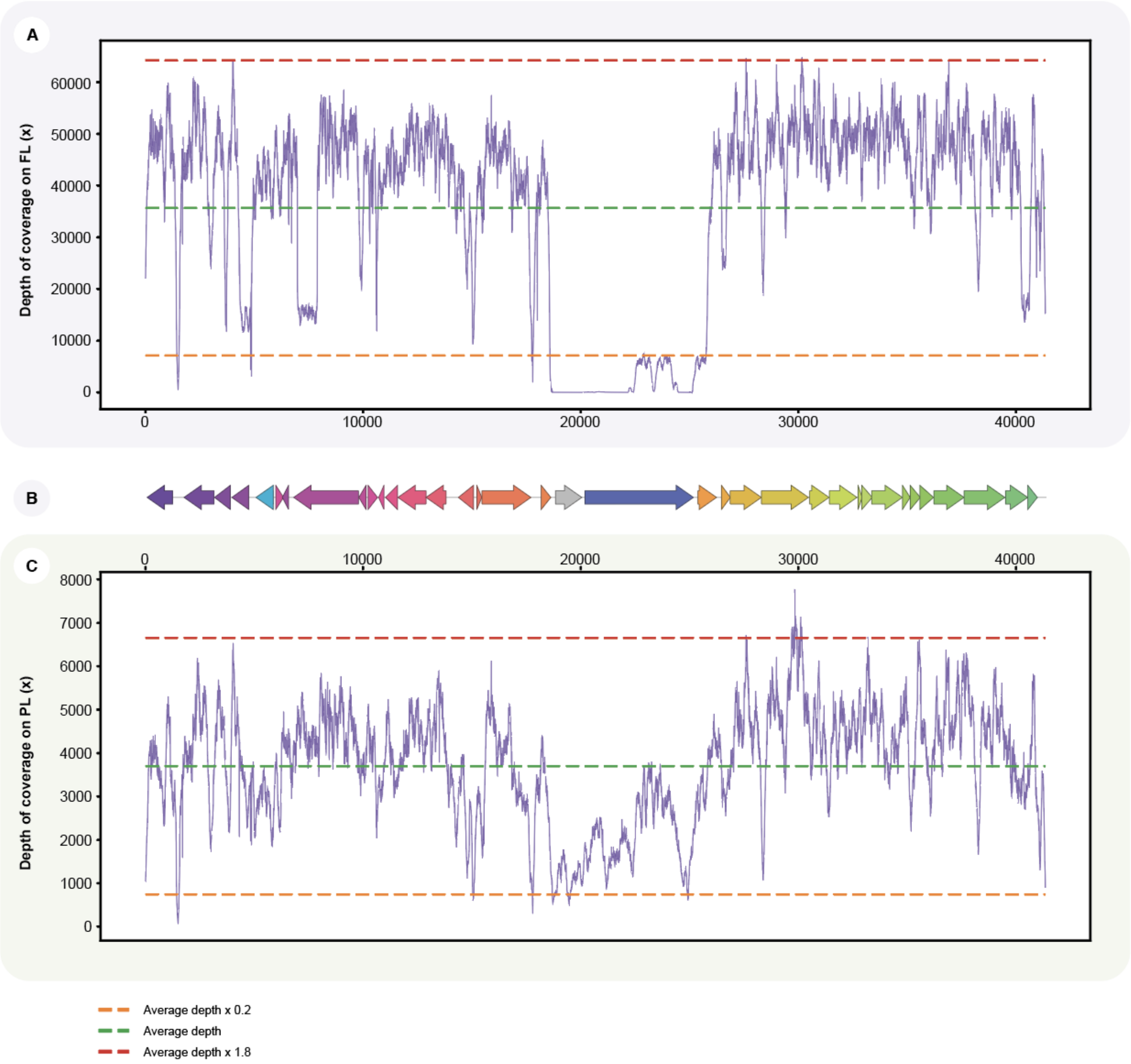


**Figure S2. The wheat-phyllosphere APSE population is differently represented in the two wheat viromes.** The read coverage of the viromes extracted from the penultimate leaf (**A**) and flag leaf (**C**) varies greatly in the region of the APSE phage genome (**B**) encoding the putative toxin cassette. The variant represented by the wheat vOTU, encoding a YD-repeat toxin (typical of the APSE-3 variants) only has coverage across the putative toxin cassette region in the flag leaf virome, revealing a great deal of variation in the toxin-encoded region across both the penultimate and flag leaf viromes.
